# Hunter-gatherers maintain assortativity in cooperation despite high-levels of residential change and mixing

**DOI:** 10.1101/313064

**Authors:** Kristopher M. Smith, Tomás Larroucau, Ibrahim A. Mabulla, Coren L. Apicella

## Abstract

Widespread cooperation is a defining feature of human societies from hunter-gatherer bands to nation states. But explaining its evolution remains a challenge. While positive assortment – of cooperators with cooperators – is recognized as a basic requirement for the evolution of cooperation, the mechanisms governing assortment are debated. Moreover, the social structure of modern hunter-gatherers, characterized by high mobility, residential mixing and low genetic relatedness, undermine assortment and add to the puzzle of how cooperation evolved. Here, we analyze four years of data (2010, 2013, 2014, 2016) tracking residence and levels of cooperation elicited from a public goods game (PG), in Hadza hunter-gatherers of Tanzania. Data were collected from 56 camps, comprising 383 unique individuals, 137 of whom we have data for two or more years. Despite significant residential mixing, we observe a robust pattern of assortment necessary for cooperation to evolve: In every year, Hadza camps exhibit high between-camp and low within-camp variation in cooperation. We further consider the role of homophily in generating this assortment. We find little evidence that cooperative behavior within individuals is stable over time or that similarity in cooperation between dyads predicts their future cohabitation. Both sets of findings are inconsistent with homophilic models that assume stable cooperative and selfish types. Consistent with social norms, culture and reciprocity theories, the data suggest that the strongest predictor of an individual’s level of cooperation in any given year is the mean cooperation of their campmates in that year. These findings underscore the adaptive nature of human cooperation – particularly its responsiveness to social contexts – as a feature important in generating the assortment necessary for cooperation to evolve.

## RESULTS

We analyze data on cooperation and residence patterns in Hadza hunter-gatherers over a six-year period (2010, 2013, 2014, 2016). Crucially, the data contain detailed information about how individual cooperative behavior persists, and how cooperators sort across time and space – vital elements that tease apart prominent theoretical models explaining the evolution of cooperation. And the presence of positive assortment of cooperators in space is a fundamental requirement of these models [1,2].

To capture the conflict between individual and group benefits we used a PG game. Games were played using an ecologically valid food item – sticks of honey. Participants could contribute 0-4 honey sticks to the PG, and all subjects split the sum of contributions multiplied by 3. Each participant’s contribution is a measure of her cooperativeness. Games were played between all adults of the same residence groups, herein called “camps”. Basic demographic information and GPS locations were recorded (Table S2). Figure 1 shows the location and mean levels of cooperation of camps in each year.

**Figure 1.**
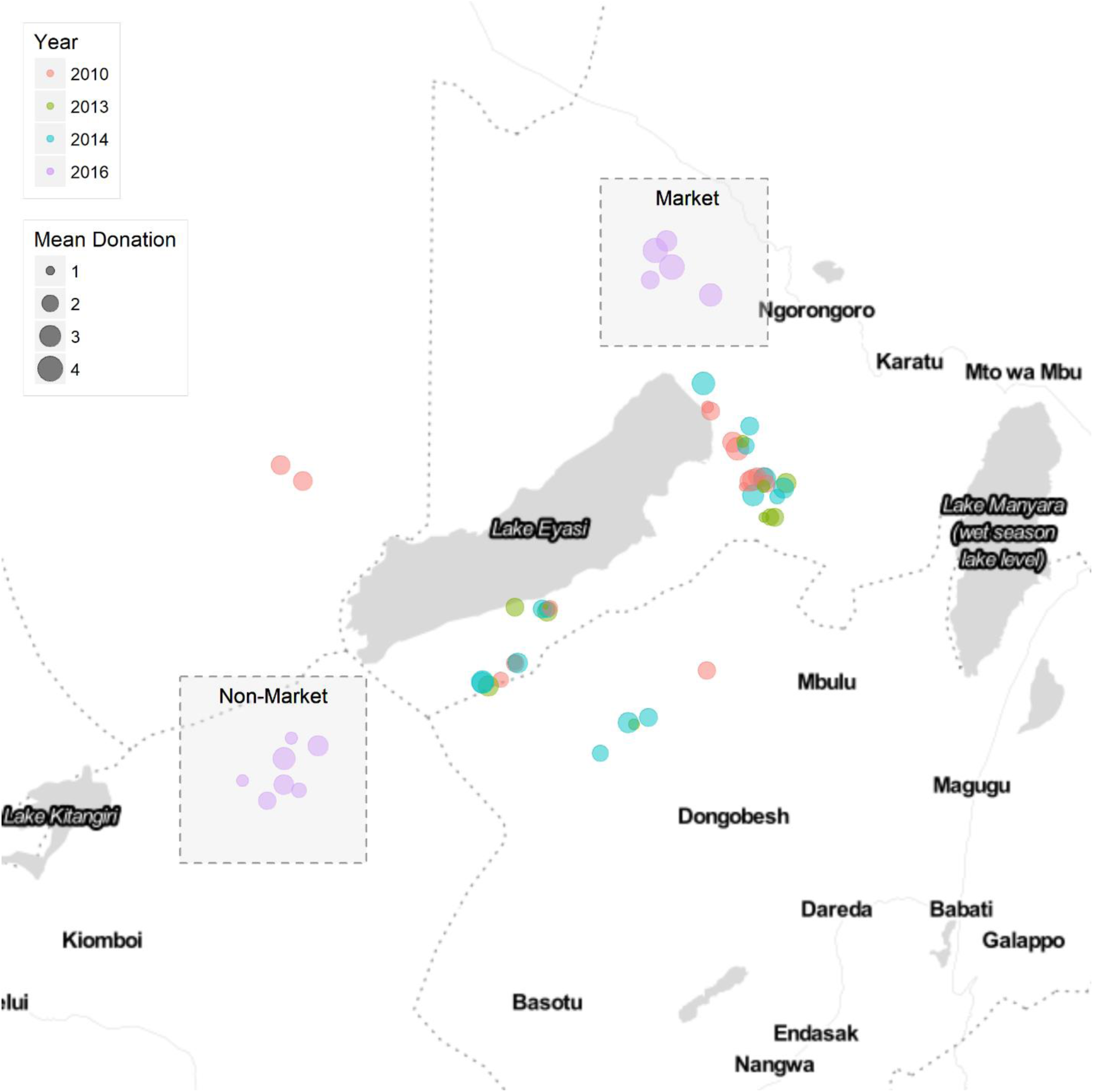
Map of the Hadza camps visited around Lake Eyasi in northern Tanzania. Circles represent the camps visited colored by year of data collection. The size of the point signifies the mean PG contribution in the camp. GPS data are not available in 2016 due to missing equipment.

### Cooperators Cluster in Camps Each Year

We first tested if individuals with similar PG game contributions cluster within camps each year. We compared the observed variance in PG game contributions with variance from 1,000 simulations. The simulations randomized participants and their contribution to different camps, but kept the population structure fixed [3]. We then measured for each simulation and the actual data the mean variance in PG contributions between participants within each camp (within-camp variance) and the variance in mean camp PG contributions across all camps (between-camp variance). In each year, less variance was observed within-camps and more variance was observed between-camps than expected in a random population (*p* < 0.05, Figure 2). The 2010 results have been previously reported [3]. The long-term data presented here suggest that assortment is a consistent feature of hunter-gatherer life, year after year. The camps in 2016 are grouped by whether they were located in the market vs non-market (see STAR method) region but their placement, is otherwise, random.

**Figure 2.**
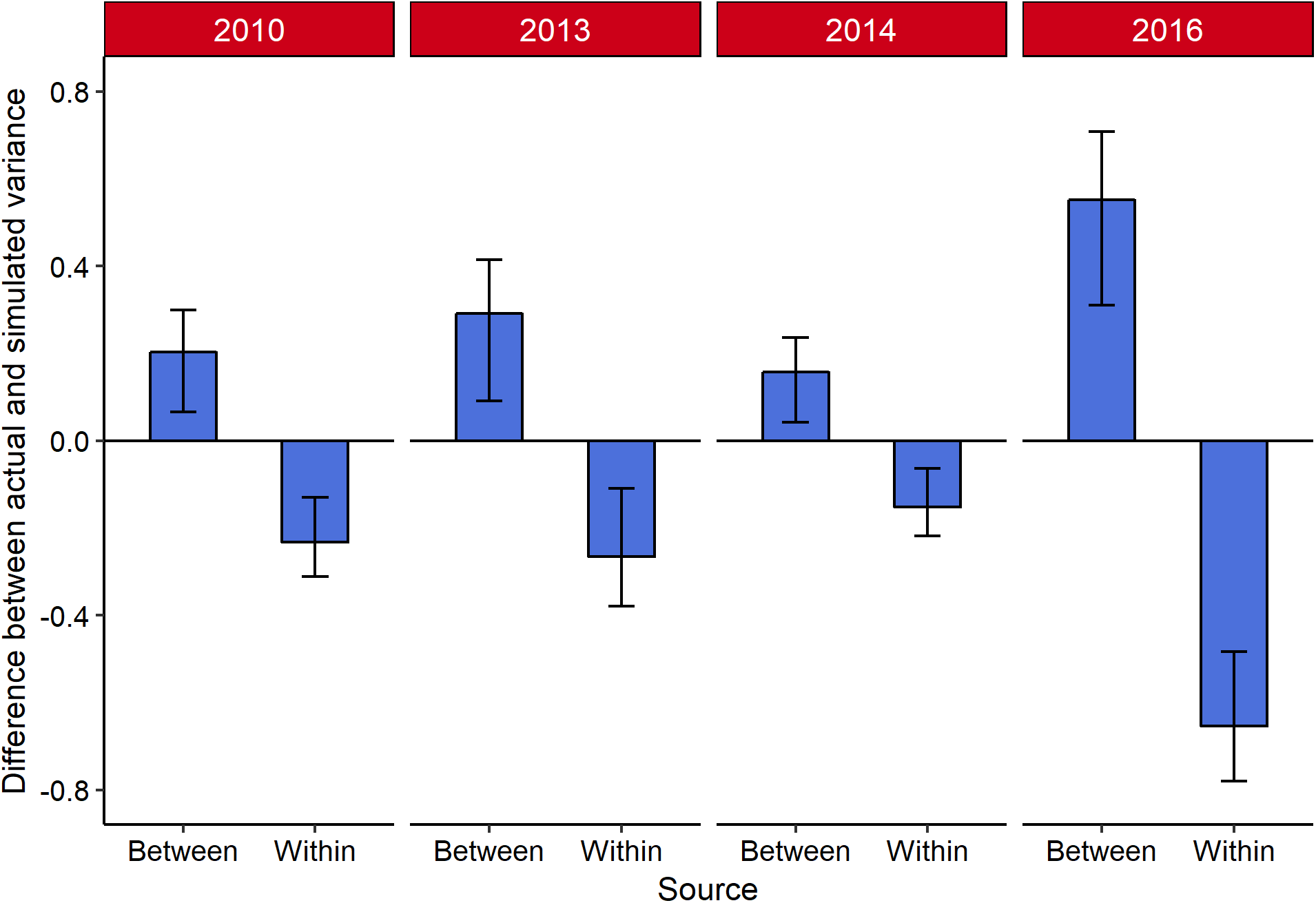
Difference between actual and simulated variance within and between residence camps in PG contributions. Error bars are 95% confidence intervals.

This observed clustering of cooperators across space meets a fundamental requirement in all theories of the evolution of cooperation: it ensures that cooperators receive benefits from other cooperators whilst remaining insulated from defectors [1]. The findings also underscore the potential for cooperation to evolve due to selection acting on groups, which may be possible when between-group variance is high relative to the within-group variance, as demonstrated in several models [4,5].

The observed assortment on cooperation in Hadza society is remarkable because the Hadza, like other hunter-gatherers, have flexible living arrangements and high rates of migration [6,7]. While common descent, where individuals preferentially interact with kin [8], and reciprocity, where individuals limit their cooperation to known reciprocators [9], can generate assortment, social mobility undermines it by 1) decreasing relatedness among group members and 2) allowing cooperative groups to be invaded by free-riders or “rovers” [10,11]. Indeed, genetic relatedness in Hadza camps is low [9].

Consistent with earlier reports [9], we too observe high rates of residential mixing in our data. We calculated for each person the proportion of campmates at time *t* that lived in same camp with the individual at time *t* + 1. The mean proportion of repeated campmates was 21.9%. Year after year, resident composition in the Hadza changes dramatically but despite this, we still see PG contributions clustering within camps each year (Figure 3).

**Figure 3.**
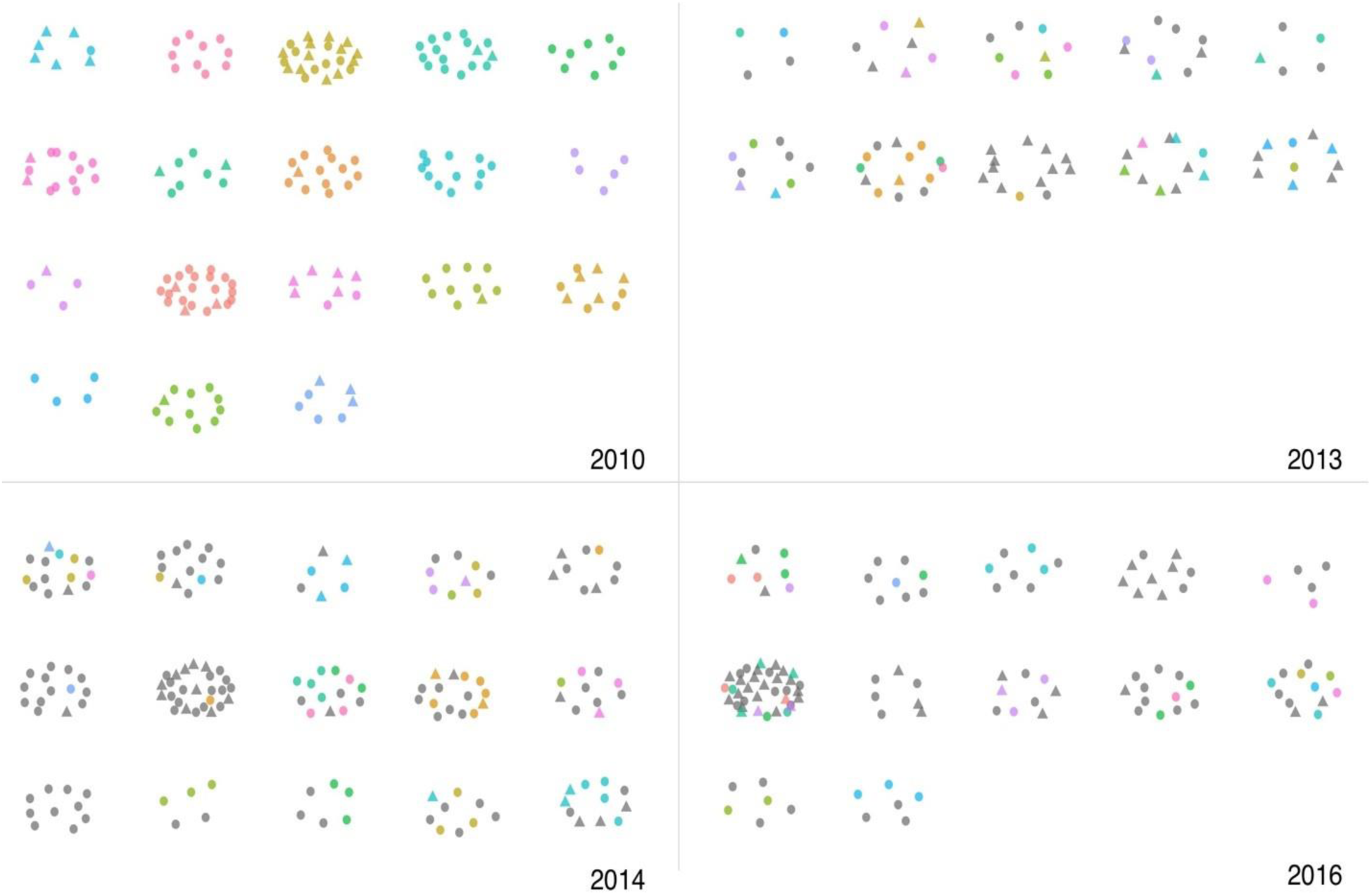
Camp residence, mixing and cooperative clustering across years. Points represent individuals grouped in space by their current camp. In 2010, individuals are colored based on current camp membership. In 2013, 2014, and 2016, individuals are colored based on camp membership in the prior wave of data collection; gray points indicate the individual was absent in the prior wave. Circles represent high cooperators (individuals who gave two or more honey sticks) and triangles represent low cooperators (individuals who gave less than two honey sticks). Camps are randomly placed in a grid.

### No Dispositional Types or Preference for Cooperators

While assortment provides an overall solution to the problem of altruism, the mechanisms responsible for real-world assortment remain unknown. One mechanism we explore is partner choice based on cooperation, in which cooperation is sustained because people choose to interact with cooperative individuals. Some of these models assume that individuals have a stable, often genetically determined, level of cooperation and individuals choose and reject partners based on this [12-14]. Under these models then, we should expect Hadza individuals to exhibit stable cooperative behavior. We should also expect that behavior in the PG at time *t* to relate to camp residency at time *t* + 1 with two possible patterns. First, if camp residency works like a market [13,15,16], with cooperative individuals being highly sought after, then we should observe individuals with similar cooperative levels at time *t* living with each other at time *t* + 1. Because cooperative individuals are highly valued, they can afford to be selective in their campmates and can choose to live with similarly cooperative individuals. This strategy is observed in human mate choice, and results in patterns of homophily [17]. Second, if camp residency does not work like a market but cooperators are still preferred, then we should observe cooperative individuals retaining more campmates between years.

We examined whether individuals’ PG contributions were related across years (Figure 4). Specifically, we examined whether current and past contributions were correlated for individuals in contiguous samples (n = 143 observations) by regressing PG contributions at time *t* on contributions at time *t* – 1 controlling for year with robust standard errors clustered on individuals and camp. There was no relationship between individuals’ current and previous contributions, *b* = 0.00, *SE* = 0.09, *t* (139) = 0.05, *p* = 0.959; this remains nonsignificant even controlling for demographic variables and exposure to markets (Table S4), and when analyzing an ordered logit regression (Table S5). It could be though that individuals prefer to give relative to the camp mean; that is, some people prefer to contribute less than, more than, or as much as their campmates across. We computed the difference between a person’s PG contribution and the mean of the rest of their campmates and repeated the analysis again with these values. Again, there was no relationship between contributions relative to campmates’ contribution at time *t* – 1 and contributions relative to campmates’ contributions at time *t, b* = 0.01, *SE* = 0.10, *t* (132) = 0.06, *p* = 0.950.

**Figure 4.**
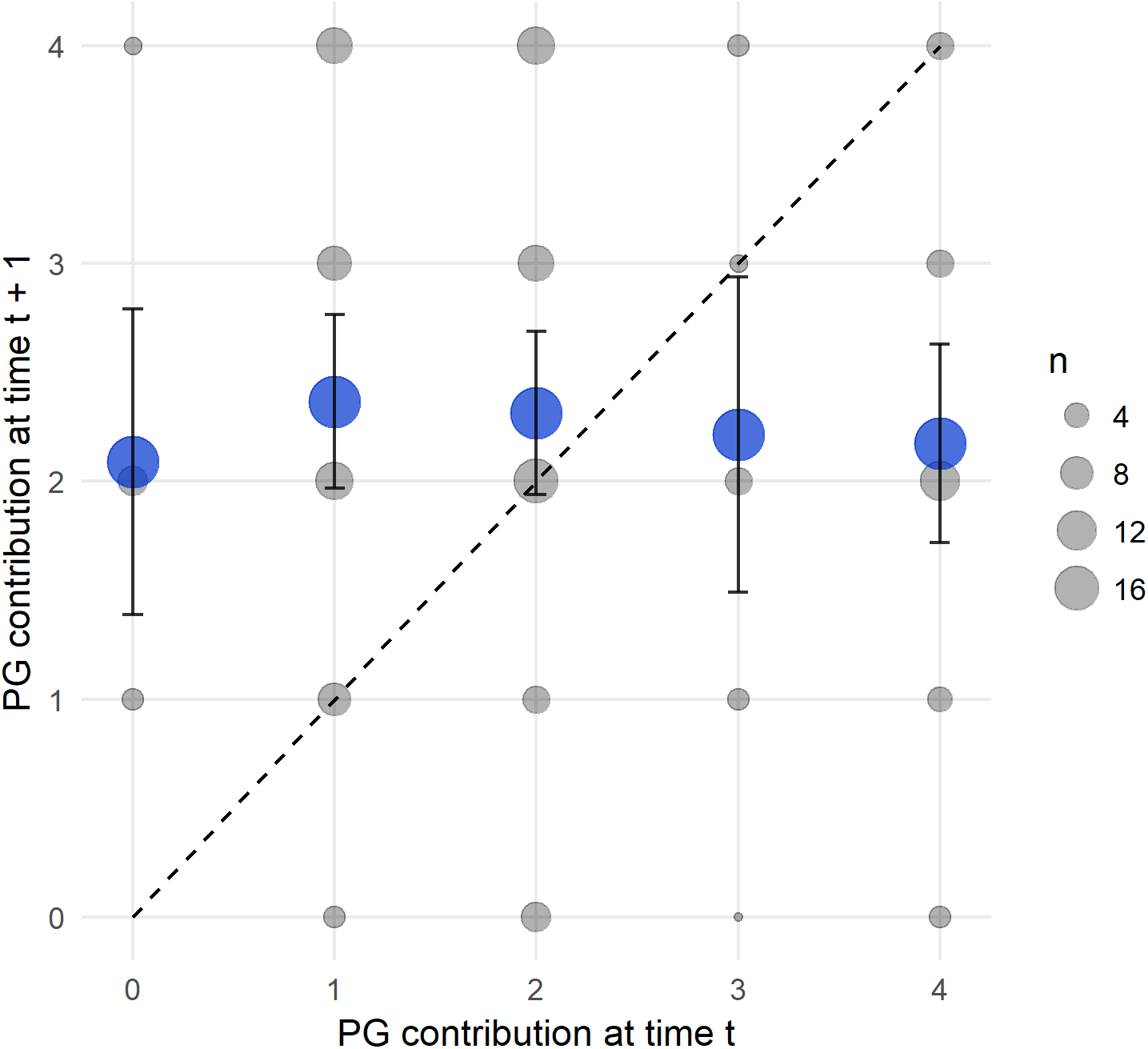
The graph plots current yearly contributions on the x-axis by contributions made in the next consecutive year on the y-axis. The unit of analysis is a participant year. Gray circles’ size is proportional to the count of individuals. Blue circles represent the average of the contribution in the following year as a function of the contribution in the current year. Bars represent 95% CI. The 45-degree line represents the null hypothesis that people have cooperative types.

We tested whether individuals with higher PG contributions were more likely to continue living with their campmates in the future. To test this, for 2010, 2013, and 2014, we calculated for each individual who was in the sample at time *t* and time *t* + 1 the proportion of campmates at time *t* that lived in same camp with the individual at time *t* + 1. We regressed PG contributions at time *t* on the proportion of repeated campmates. Robust standard errors were clustered on the individual and the camp. There was, in fact, a negative, but nonsignificant, relationship. Individuals who contributed more at time *t* had fewer repeated campmates at time *t* + 1, *b* = – 0.02, *SE* = 0.01, *t* (141) = -1.92, *p* = 0.057. It could be that cooperative individuals are leaving their previous campmates and being selective with their new campmates, choosing to live with similarly highly cooperative individuals. To test this, we regressed the mean contribution of campmates at time *t* + 1 on individuals’ contribution at time *t* with robust standard errors clustered on the individual. There was no relationship between a person’s contribution in one year and the mean contribution of their campmates the next year, *b* = −0.03, *SE* = 0.05, *t* (141) = – 0.57, *p* = 0.568. Cooperative individuals were not more likely to continue living with their campmates or to find more cooperative campmates.

To further test if cooperative individuals were choosing to live with similarly cooperative individuals, we tested if similarity in PG contributions in a past year predicted whether Hadza will live together in a future year. We created a dataset for 2010, 2013, and 2014 of every possible dyad in each year, removing dyads if neither individual was present in the next sample. This resulted in 21,086 observations with 18,126 unique dyads across years. Of these observations, 789 (3.9%) of dyads were in the same camp. Using a binary logistic regression, we regressed whether the dyad lived in the same camp at time *t* + 1 on the similarity of PG contributions at time *t* and whether the dyad lived in the same camp at time *t* with robust standard errors clustered on the dyads. Individuals who contributed similar amounts were not more likely to live in the same camp in future years, *b* = 0.01, *SE* = 0.04, *OR* = 1.01, *Z* = 0.24, *p* = 0.814, which remained nonsignificant after controlling for demographics variables (Table S3).

### Campmates Influence Cooperative Behavior

To explore the role of social context on cooperative behavior we tested whether an ego’s contribution can be predicted by the mean contribution of their current campmates, controlling for various demographic variables. First, we calculated for each person a camp mean contribution excluding ego’s own contribution. We regressed PG contributions of ego on the mean contribution of other camp members controlling for year with errors clustered on the individual and camp. Corroborating the analyses simulating between and within-camp variation, we find that for each additional honey stick contributed by camp members, ego contributed on average another half-stick of honey, *b* = 0.55, *SE* = 0.15, *t* (138) = 3.60*, p* < 0.001. This remained significant when controlling for various demographic variables (Table S4) and when analyzed using an ordered logit regression (Table S5). Further, in 2010 and 2016, the only years for which we have kinship data (see STAR methods), we regressed PG contributions on campmates’ mean contributions controlling for close relationships with robust standard errors clustered on the individual and camp. Campmates’ mean contributions remained significant in this regression, *b* = 0.79, *SE* = 0.06, *t* (314) = 12.53, *p* < 0.001.

For participants in which we have overlapping data across years, we also examine whether the mean contribution of an ego’s current campmates is a better predictor of ego’s current contribution, than ego’s own past contribution. For each year, we regressed ego’s current contribution on the mean contribution of their current campmates and ego’s contribution in the previous year with robust standard errors clustered on the individual and camp. For each additional honey stick given by camp members, ego again contributed an additional half-stick of honey, *b* = 0.50, *SE* = 0.16, *t* (132) = 3.11*, p* = 0.002. There was still no effect of previous contribution on current contribution, *b* = -0.01, *SE* = 0.08, *t* (132) = -0.15, *p* = 0.879. The results did not change when controlling for demographic variables (Table S4) or when using an ordered logit (Table S5). We also find no evidence that having played the game in a prior year predicts subsequent contributions (Table S6).

## DISCUSSION

While multiple theoretical models to explain the evolution of cooperation have been proposed, there is little evidence on what theories actually explain cooperation in evolutionarily-relevant settings. The Hadza provide an important test case for evolutionary models of cooperation: Their daily life is marked by widespread sharing of food, labor, and childcare. And their lifeways more closely approximate pre-Neolithic populations compared to samples drawn from Western, Educated, Industrialized, Rich and Democratic (WEIRD) societies [18].

We show that positive phenotypic assortment in camp residence is a characteristic feature of Hadza society year after year. This structuring of the population satisfies a basic requirement underlying all theories for the evolution of cooperation – cooperators must cluster together in social space [1,2]. The significance of this finding is further underscored by the fact that the Hadza exhibit high levels of mobility and inter-group mixing – features that undermine assortment in classical models of cooperation. Mobility provides defectors opportunities to invade cooperative groups [10,11] and lowers relatedness within groups [19].

For this reason, additional classes of theoretical models explaining how assortment on cooperation can be achieved have been emphasized. In models of biological markets involving partner choice, individuals compete for the most cooperative partners and the most cooperative choose each other [16]. In models involving conditional strategies that respond to group-level behaviors, such as generalized reciprocity [20] and/or the switching of groups [21], cooperation can stabilize when the groups are small [20]. In models of gene-culture co-evolution, culturally evolving social norms, supported by an underlying norm-psychology, can generate within-group similarity and between-group differences in cooperation [22].

While nearly all models involve some degree of behavioral flexibility such that an individual’s level of cooperation is contingent on the social environment, some partner choice models assume that individuals have stable dispositions on which the choice of partners is based [12-14,21]. And, individuals can leave current partners or reject prospective partners based on their observations and past interactions. In some models, the market generates a selection pressure for cooperation, leading to the genetic evolution of the trait.

The real-world application of these models hinges on the existence of stable differences in cooperativeness. We find no evidence that cooperative behavior is stable over time. We also do not find that similarity in cooperation between individuals predicts their future co-habitation or that cooperators retain more campmates over time.

Natural selection should favor individuals who select partners based on how much they actually cooperate, which is determined by both their partner’s willingness and ability to cooperate [16]. Whether willingness or ability to cooperate is valued more as a criterion for partner choice will depend, in part, on which trait is more variable in the population [23]. In laboratory studies, participants do display a preference for partners who are willing to cooperate, possibly because cooperative contributions are artificially constrained in these settings. Conversely, the Hadza have strong norms governing cooperation and sharing. If everyone shares because they are expected to, then one’s ability to share may be valued more than their willingness to share. In fact, prior interview data suggest that when given the choice, the Hadza do not choose the most cooperative individuals as campmates [3]. Instead physical traits show a small, but positive correlation with how often individuals are chosen as campmates, possibly because provide some indication about their ability to acquire resources. Whether the Hadza trade-off willingness to cooperate for other qualities, such as foraging ability, and/or whether partner choice involves some degree of complementarity where individuals seek partners that provide different types of benefits [24] would be an interesting avenue for future study.

Few studies have examined longitudinal stability in cooperation. Our findings contrast with laboratory studies using Western samples illustrating small-to-medium-sized correlations in cooperative game play over time [25,26]. The discrepant results may also be due to the longer intervals between testing in our study. Also, the Hadza are playing the game with different, but well-known, individuals each year. In laboratory settings, individuals are often playing in the same anonymous/unfamiliar group setting each time. Finally, cultural differences in dispositional consistency may also explain the divergent results. Compared to individuals from collectivist societies, Westerners tend to describe themselves in terms of underlying psychological traits and have a stronger preference for self-consistency [27]. Interestingly, in a small sample (*n* = 12) of Tsimane' forager-horticulturalists, no relationship was found in dictator game play measured two years apart [28].

While we cannot isolate the exact mechanism(s) generating the within-group homogeneity on cooperation, we find that cooperative behavior in any given year is best predicted by the cooperativeness of one’s current residence group, not demographics, number of kin in one’s camp or past cooperative behavior. The results are consistent with social learning and reciprocity theories of cooperation and concur with laboratory experiments demonstrating that cooperative and selfish play in economic games is contagious [29,30].

By using an economic game as our measure of cooperation, as opposed to measuring naturally occurring levels of cooperation, we traded-off some ecological validity for increased experimental control. We chose the PG game due to its direct relevance to hunter-gatherer life where collective action problems are a daily occurrence. We observe that across years, the Hadza, on average, contribute 56% of their endowment to the PG. These results provide some reassurance that local institutions are, in fact, mapping onto game play.

While we attempted to create maximal experimenter control, it is difficult to establish the same degree of control in field settings that are found in the laboratory. Thus, the problem of omitted variable bias is a genuine concern as there may be other influences on cooperation that were unobserved. Future work would benefit from more in-depth examinations into other factors that influence Hadza decisions to cooperate, possibly through interviews and experiments.

A third limitation of the study is that we collected data at discrete points far apart in time. As a result, we are limited by how much we can say about the formation and breakdown of camps in relation to cooperation. Hunter-gatherer residence is determined by multiple and complex demographic, economic, ecological and personal factors [3,7] – all phenomena outside the purview of the current study. Future work examining the role of cooperation in Hadza camp formation and dissolution would be an important area for future exploration.

Finally, we suggest that studying the conduits of norm establishment and reinforcement in hunter-gatherers hold particular promise. Storytelling, for instance, may be an effective way to teach and establish norms [31]. Recently, it has been documented that among Agta foragers, groups with more skilled storytellers are more cooperative [31]. Moreover, there is a large literature demonstrating how ritual activities, which are thought to enable the expression of shared beliefs and norms, can impact cooperation and fairness [32]. Hadza life is replete with important public and private ritualistic activities – including song, dance, meat-eating, storytelling and puberty initiation practices – which are thought to play an important role in cementing relationships and promoting cooperation [7].

In summary, we find that the best predictor of an individuals’ level of cooperation is the mean cooperativeness of their current social group. The findings are consistent with evolutionary models stressing the importance of contingent reciprocity, cultural learning and social norms [22,33,34] and highlight the remarkable capacity of humans to respond adaptively to their social environments.

## STAR METHOD

### Contact for resource sharing

Further information and requests for resources and reagents should be directed to and will be fulfilled by the Lead Contact, Coren Apicella

### Subject details

#### Study Site

The Hadza are nomadic foragers occupying the Lake Eyasi basin within the Great Rift Valley in Northern Tanzania. They sleep outside under the stars or in makeshift huts constructed of grass and trees. Approximately 1,000 individuals identify as Hadza, but only 200-300 individuals obtain the majority of their calories by hunting and gathering. It is this latter group that is the focus of this research.

Men hunt birds and mammals using bows and poison-tipped arrows and collect honey. Women gather plant foods including baobab fruit, berries, and tubers. Food is shared widely within camps, especially big game but producers of the food can channel the food in ways that benefit their kin [35]. Childcare is also shared [36].

The Hadza live in temporary camps that average about 30 individuals. Camps generally consist of several unrelated nuclear families. Relatedness within camps is low with primary kin comprising, on average, 1.43 and 1.93 of men and women’s campmates respectively [7]. Typical of most contemporary hunter-gatherers, residence patterns are fluid and are best described as fission-fusion grouping [37]. Camps can merge or split. Individuals too, can freely relocate to new camps. Every 4-8 weeks entire camps shift location usually in response to resource availability. Because the Hadza have few capital goods and personal possessions, the physical costs associated with moving remain low.

While there is striking diversity among forager societies, it is thought that the social, economic and political arrangements of the Hadza are similar to other hunter-gatherer societies. A study of hunter-gatherer social life using ethnographic data from 437 past and present foraging societies found that the vast majority of forager societies, including the Hadza, live in small groups, practice central place foraging and food sharing [37]. The Hadza also fall at or near the median value on a variety of key demographic traits such as the percentage of calories contributed to the diet by men and women, infant mortality rate, fertility rate, inter-birth intervals and so on [37]. Thus, apart from the fact the Hadza still maintain a subsistence lifestyle, there is good reason to believe that they are not outliers in other major respects.

Ethno-tourism, which largely began about 10-15 years ago has had the largest impact on Hadza life. And tourists visiting the Hadza continue to rise each year. While tourists can now be found in every region of Hadzaland, the vast majority of visits take place in camps on the north-eastern side of Lake Eyasi, close to the village Mangola, due to its proximity to paved roads that lead to Arusha and safari parks (Figure 1). Tours usually last a couple of hours and culminate with a cash payment to the camp which then the Hadza can spend in the village.

The Hadza have been described as having little belief in omniscient, moralizing gods [37,38] but they do engage in a number of important rituals including a sacred epeme dance and meat-eating rituals [37]. These rituals are thought to bond participants to one another [7].

#### Sample Characteristics

Across years, we visited 56 Hadza camps collecting data from 383 unique individuals. For 137 participants, we have data from at least two years (Table S1). The mean age was similar across the years, ranging from 37 to 40 and women comprised 51%, 42%, 49% and 46% of the sample in 2010, 2013, 2014, and 2016, respectively. Further summary statistics can be found in the supplementary materials (Table S2).

#### Ethical Permissions

Institutional approvals were obtained prior to conducting this study from the Committee on the Use of Human Subjects at Harvard University, The University of Pennsylvania Institutional Review Board and the Tanzanian Commission for Science and Technology (COSTECH). Verbal informed consent was obtained from all participants due to low literacy rates.

### Method details

#### Data Collection

Data was collected in four separate years – usually during the dry season – over a six-year period (2010, Aug/Sept; 2013, July; 2014, Oct/Nov; 2016, Aug/Sept). Data collection was supervised by different authors in different years: (CLA in 2010, 2013; IM in 2014 and KMS in 2016). In 2014 and 2016 Tanzanian researchers blind to the hypotheses collected the data. In each year, camps were visited using a technique not unlike snowball sampling. After establishing contact with the first camp, Hadza would direct the researchers to the next nearest camp. GPS coordinates were recorded for all camps in each year, with the exception of 2016 when the GPS receiver met an unfortunate end. Nevertheless, we were able to divide the camps in 2016 into market and nonmarket groups based on their general proximity to the village (Figure 1).

#### Public Goods Game

We used a public goods game as our measure of cooperation. This game is directly applicable to hunter-gatherer life where collective action problems are faced by groups on a daily basis. We used a food item instead of money since explanations for the evolution of cooperation have highlighted the importance of food sharing [39-41]. The methods for the PG game elicitation in the Hadza has been described previously [3].

Cooperation was elicited by examining participants’ voluntary contributions in a public goods game played with adult members of their camp. All games were conducted in Swahili and inside a vehicle for privacy. All adults in each camp were invited to participate with the exception of the very elderly and infirm. In 2010, 2013 and 2014 the game was played on the last day the researcher was in camp in order to limit possible discussion. Participants were also told that the game was secret. Since decisions were made in private, any assertions made by participants regarding their decision need not be truthful. In 2016, the game was played throughout the researcher’s stay in the camp. Importantly, we find the same pattern of results.

Participants were endowed with four straws of 100% pure honey (2010, Honeystix, GloryBee foods Inc. 2013, 2014, Honey Stix, Stakich Inc.), a prized food of the Hadza [42]. Each honey stick contains roughly 15 calories. Participants then faced the decision of how to divide their honey sticks into a private account and a public account. Participants were told that the goods would be distributed evenly with all other adult camp members who also played the game. They were instructed that they could keep any amount from 0-4 sticks of the honey or donate them to the public good by inserting them into an opaque cardboard box with an opening at the top. Subjects were told that for every stick of honey they donated, the researcher, would donate an additional 3 sticks of honey to the public pot, and that, after all adult campmates played the game, the honey would be divided equally among them. Participants were also told that they would receive their undonated honey at the same time as the public honey was distributed to avoid confounding generosity with patience. Participants were also told that the game was a secret. Before subjects made their decision, the researcher simulated all their possible choices so that subjects were shown the additional amount of honey added to the box for each decision.

The Hadza have had experience playing various games to measure economic (e.g. endowment effect and risk) and social preferences (e.g. dictator, ultimatum, third-party punishment) with researchers over the last decade [43-46].

This is the basic script used each year in both English and Swahili.

### English

We are playing a game with honey. This game is voluntary. You do not have to play this game. You will not be punished if you choose not to play. This study is a secret. I will not tell anyone the decision you make. Also, I will not tell you the decision that anyone else makes. All adults living in your camp will have the opportunity to play this game.

This game involves honey (*show them 4 honey sticks*). Inside these sticks is honey to eat. The decisions you make and the decisions other people make will affect how much honey you get and how much honey your other camp members get. You will only receive your share of honey after everyone has had a chance to the play the game. Any honey you receive will be given to you in secret, and nobody will see how much honey you get.

Here are 4 sticks of honey (*hand it to them*). You need to choose how many sticks to keep and how many sticks to put inside this box. You can choose to:

keep all of the sticks of honey

keep 3 of the sticks

keep 2 of the sticks

keep1 stick

keep zero sticks.

No one will know how many sticks you choose to keep. Any honey that you do not keep will be put in this box and shared equally with all the people who played this game, including yourself. For every stick of honey you put in this box, I will add 3 sticks.

If you put in 1 stick, I will add 3 sticks.

If you put in 2 sticks, I will add 6 sticks.

If you put in 3 sticks, I will add 9 sticks.

If you put in 4 sticks, I will add 12 sticks.

If you keep all 4 honey sticks for yourself, I will not add any honey to the box.

If everyone puts honey in the box, then the box will fill up and everyone will get a lot more honey. If no one or only a few people put honey in the box, then there will be very little honey to share.

### Swahili

Tunaenda kucheza mchezo wa asali. Mchezo huu ni hiari. Unaweza kuamua usicheze mchezo huu. Hautaadhibiwa kama utaamua kutocheza. Somo hili ni siri. Sitamwambia mtu yeyote maamuzi utakayofanya. Pia, sitakwambia maamuzi ambayo mwingine amefanya. Watu wazima wote wanaoishi kwenye kambi yako watakuwa na nafasi ya kucheza mchezo huu. Mchezo huu unahusisha asali (waoneshe fimbo 4 za asali). Ndani ya fimbo hizi ni asali unaweza kuila. Maamuzi ambayo unafanya na maamuzi ambayo watu wengine wanafanya yanaathiri jinsi wewe unavyopata asali na watu wengine pia kambini. Utapata tu sehemu yako ya asali baada ya kila mtu kupata nafasi ya kucheza mchezo. Na asali utakayopata utapewa kwa siri na hakuna yeyote atakayeona umepata asali ngapi.

Hizi ni fimbo 4 za asali (mkabidhi). Unatakiwa uchague ni fimbo asali ngapi ubakiwe nazo na asali ngapi uweke ndani ya boksi hili. Unaweza kuchagua:

Kubakiwa na fimbo zote za asali

Kubakiwa na fimbo 3

Kubakiwa na fimbo 2

Kubakiwa na fimbo 1

Kutobakiwa na fimbo, sifuri

Hakuna mtu ambaye atajua umeamua kubakiwa na fimbo ngapi

Na asali yeyote ambayo hutobakiwa nayo itawekwa ndani ya boksi hili na zitagawanywa sawa kwa sawa na kila mtu ambaye amecheza mchezo huu, ukiwemo wewe

Kwa kila fimbo ya asali utakayoweka ndani ya boksi hili, nitaongeza fimbo 3.

Ukiweka fimbo 1, nitaongeza fimbo 3

Ukiweka fimbo 2, nitaongeza fimbo 6

Ukiweka fimbo 3, nitaongeza fimbo 9

Ukiweka fimbo 4, nitaongeza fimbo 12

Kama utabakiwa na fimbo zote 4 za asali kwa ajili yako, sitaongeza asali yeyote ndani ya boksi

Kama kila mtu ataweka asali kwenye boksi, hivyo boksi litajaa na kila mtu atapata asali nyingi sana. Kama hakuna mtu au watu wachache wataweka asali kwenye boksi, kutakuwa na asali kidogo sana za kugawana/shirikiana

#### Additional Control Variables

*Basic Demographics*. Age, marital status, spouse’s names and reproductive histories were recorded each year.

*Education*. Participants were asked the number of years that they attended school in 2013 and 2016. PG contributions were regressed on the number of years of formal education.

*Household size*. We asked participants the number of other individuals living in their household in 2013 and 2016. This typically includes children and spouse and occasionally other close family members. We regressed PG contributions on household size.

*Concerns about food*. In 2013, participants were asked two forced choice questions about whether they were worried there would be enough food for their family in 1) over the next month or 2) over the year. Participants answered yes or no to both questions, such that a “yes” indicated participants were worried about having enough food.

*Trade*. In 2013, participants were asked to estimate how many days out of the past seven they personally went to a market or trade center to buy or sell something.

*Close Relationships in Camp*. In 2010 and 2016, we asked participants to provide the names of their biological parents, which allowed us to identify primary kin (full siblingships and parent-child relationships) living together. For each individual, we then calculated the proportion of their campmates that were primary kin or a spouse as a measure of “close relationships.”

*Time of Day*. In 2010, 2013, and 2014, the public good game was played after all other data were collected and in a short time period. Time was not recorded in these three sample years. In 2016, the public good game was played throughout the study period so that the time the game was played varied within camps. Time of day was categorized into three periods: morning if the game was played between 8:00 and 12:00, afternoon if played between 12:00 and 16:00, and evening if played between 16:00 and 18:00.

### Quantification and Statistical Analysis

#### Software

All analyses were conducted in R. For data manipulation, we used the tidyverse [47], magrittr [48], and dplyr [49] packages. For regression analyses with robust standard errors, we used the lmtest [50], multiwayvcov [51] and sandwich [52] packages. For visualizations, we used the ggplot2 [53], scales [54], gridExtra [55], GGally [56], RColorBrewer [57], ggmap [58], geosphere [59], network [60], sna [61], and igraph [62] packages.

#### Variance in PG contributions

To test if PG contributions clustered within camps, we measured variance between camps and variance within camps in PG contributions. Variance between camps was the variance in camp mean contributions between camps, and variance within camps was the mean variance within each camp between individuals in PG contributions. For each year, we then simulated the population distribution of these values. PG contributions were randomly re-assigned without replacement within the population structure. For each run, the variance between and within camps in PG contributions was saved. The actual variances were compared to the distribution of simulated variances; if the actual variances fell within the extreme tales of the distribution (2.5% or 97.5%) the variances were determined to be significantly different from chance.

#### Regression analyses

For regression analyses that did not involve variables from previous years, all observations in 2010, 2013, 2014, and 2016 were used. All models had robust standard errors clustered on the individual. For models that include mean camp PG contribution, we calculated for everyone the mean of other camp members’ contribution such that an individual’s mean camp PG contribution did not include ego’s own contribution. For these analyses, robust standard errors were also clustered on the camp. For regression analyses that involved variables from previous years, observations in 2013, 2014, and 2016 were included only if the individual was in the previous sample year. For these analyses, robust standard errors were clustered on the individual, and if the analysis include mean camp PG contribution, they were clustered on the camp as well.

Given the limited range possible in PG contributions, it could be argued that these data should be analyzed as if they were ordinal. Here, we re-run the key analysis regressing individual PG contributions on mean camp contributions and previous contributions using an ordered logit to test the robustness of our results. Again, we limit the analysis to contributions in 2013, 2014, and 2016 including only participants who also had contributions in the previous year. Again, we clustered the robust standard errors on the individual and camps.

#### Analysis of Dyads Living Together in Future Years

We constructed a dataset of dyads to analyze who lives with whom in each year. To do this, we went through 2010, 2013, and 2014 and for each individual *i* in the sample at time *t* and time *t* + 1, we went through each individual *j* at time *t* and recorded whether *i* and *j* lived in the same camp at time *t*, at time *t* + 1, and their similarity in PG contributions at time t, as well as their similarity on demographic variables at time t. Similarity scores were calculated by finding the absolute value of the difference between *i* and *j* on the variable and multiplying that value by -1 so that greater values indicate more similarity on the variable. We used a binary logistic regression and regressed whether *i* and *j* lived together at time *t* + 1 on the other variables with robust standard errors clustered on dyads.

## Author contributions

CLA and KMS contributed to study design and writing. CLA, KMS, TL analyzed the data. CLA, IAM, KMS collected data. All authors commented on drafts of the manuscript.

The authors declare no competing financial interests.

---

**Table S1.** Sample Sizes Within and Across Years. Related to STAR methods

| Year | 2010 | 2013 | 2014 | 2016 |
| --- | --- | --- | --- | --- |
| 2010 | 191 | 46 | 69 | 42 |
| 2013 |  | 99 | 57 | 31 |
| 2014 |  |  | 170 | 40 |
| 2016 |  |  |  | 127 |

*Note.* Total number of participants in each year on the diagonal. Other cells indicate number of participants in both years.

**Table S2.** Demographic Variables and Their Relationship to PG Contributions. Related to STAR Method and Results.

| Measure | 2010 | 2013 | 2014 | 2016 | Relationship<br>with PG<br>contributions |
| --- | --- | --- | --- | --- | --- |
| PG contributions | 2.3 (1.2) | 1.7 (1.2) | 2.5 (1.1) | 2.2 (1.4) |  |
| Males | *n* = 94 | *n* = 57 | *n* = 86 | *n* = 58 | 0.10 (0.10) |
| Married | *n* = 152 | *n* = 76 | *n* = 130 | *n* = 90 | -0.06 (0.13) |
| Age | 37.1 (11.0) | 40.0 (12.9) | 39.6 (13.4) | 37.6 (14.6) | 0.01 (0.004) |
| Number of living children | 3.1 (2.3) | 3.3 (2.4) | 3.5 (2.6) | 3.2 (2.6) | -0.02 (0.03) |
| Close relationships | 0.12 (0.12) |  |  | 0.14 (0.16) | -1.11 (2.34) |
| Formal education |  | 1.4 (2.7) |  | 1.2 (2.5) | 0.01 (0.06) |
| Household size |  | 4.2 (2.2) |  | 2.7 (2.0) | 0.00 (0.06) |
| Food concern for the next month |  | *n* = 56 |  |  | -0.74 (0.44) |
| Food concern for the next year |  | *n* = 53 |  |  | -0.51 (0.53) |
| Trade |  | 0.5 (0.8) |  |  | 0.15 (0.19) |

*Note.* For descriptive statistics, values are counts or mean (standard deviation in parentheses) for that variable in each year. See STAR method for more description of each variable. Values in the “Relationship with PG contributions” are unstandardized coefficients (standard error in parentheses) of contributions regressed on that variable only—that is, each variable was entered into a separate regression—with robust standard errors clustered on the individual and camp. None of the variables significantly related to PG contributions.

**Table S3.** Binary Logistic Regression on Dyads Living in the Same Camp. Related to No Dispositional Types or Preference for Cooperators

|  | *b* (*SE*) | *OR* | *Z* | *p* |
| --- | --- | --- | --- | --- |
| Intercept | -3.51 (0.17) | 0.03 | -20.37 | < 0.001 |
| Lived together previously | 0.37 (0.14) | 1.44 | 2.56 | 0.010 |
| Similarity in PG contributions | 0.01 (0.04) | 1.01 | 0.24 | 0.814 |
| Both male | 0.18 (0.11) | 1.20 | 1.71 | 0.087 |
| Both female | 0.28 (0.10) | 1.33 | 2.74 | 0.006 |
| Both married | -0.01 (0.09) | 0.99 | -0.10 | 0.922 |
| Both single | -0.67 (0.33) | 0.51 | -2.03 | 0.042 |
| Similarity in age | 0.01 (0.004) | 1.01 | 1.65 | 0.099 |
| Similarity in number of living children | 0.05 (0.02) | 1.05 | 2.47 | 0.014 |
| Both lived in market region previously | 0.13 (0.11) | 1.13 | 1.10 | 0.273 |
| Both lived in non-market region previously | 0.48 (0.10) | 1.62 | 4.75 | < 0.001 |

*Note.* Whether the dyad lived in the same camp at time $t + 1$ was regressed on variables in the model. All variables in the model are taken from time $t$ . See STAR method for more detail on the analysis.

**Table S4.** OLS Regressions of PG Contribution on Mean Camp Contribution and Previous Contributions. Related to Campmates Influence Cooperative Behavior.

|  | Model 1 | Model 2 | Model 3 | Model 4 | Model 5 |
| --- | --- | --- | --- | --- | --- |
| Mean camp contribution | 0.55***<br>(0.15) | 0.36*<br>(0.16) |  |  | 0.36*<br>(0.16) |
| Previous contribution |  |  | 0.00<br>(0.09) | -0.01<br>(0.08) | -0.01<br>(0.08) |
| 2014 | 0.44*<br>(0.19) | 0.53*<br>(0.22) | 0.75**<br>(0.24) | 0.76**<br>(0.25) | 0.53*<br>(0.23) |
| 2016 | 0.50<br>(0.26) | 0.76**<br>(0.23) | 0.76<br>(0.39) | 1.05***<br>(0.23) | 0.76**<br>(0.23) |
| Male |  | 0.17<br>(0.19) |  | 0.18<br>(0.19) | 0.17<br>(0.19) |
| Age |  | 0.00<br>(0.01) |  | 0.00<br>(0.01) | 0.00<br>(0.01) |
| Married |  | 0.25<br>(0.31) |  | 0.33<br>(0.29) | 0.25<br>(0.31) |
| Number of living children |  | -0.03<br>(0.03) |  | -0.04<br>(0.03) | -0.03<br>(0.03) |
| Exposure to market |  | -0.03<br>(0.19) |  | -0.04<br>(0.25) | -0.03<br>(0.20) |
| Coresidents at time *t* |  | -0.03*<br>(0.01) |  | -0.05***<br>(0.01) | -0.03*<br>(0.01) |

*Note.* Values are unstandardized OLS regression coefficients with standard errors in parentheses. Coresidents is the number of other Hadza who played in the PG game. The first two models test the effect of campmates' contributions on ego's contributions, with the second model adding demographic controls. The third and fourth model test to what extent current contributions are associated with previous contributions, with the fourth model adding demographic controls. Finally, the fifth model tests both effects together, with the demographic controls. All analyses are restricted to contributions in 2013, 2014, and 2016, and to individuals with a previous contribution in the sample year prior.
\* $p < 0.05$ , \*\* $p < 0.01$ , \*\*\* $p < 0.001$

**Table S5.** Ordered Logit Regressions of PG Contribution on Mean Camp Contribution and Previous Contributions. Related to Campmates Influence Cooperative Behavior.

|  | Model 1 | Model 2 | Model 3 | Model 4 | Model 5 |
| --- | --- | --- | --- | --- | --- |
| Mean camp contribution | 0.96***<br>(0.28) | 0.69**<br>(0.31) |  |  | 0.69**<br>(0.31) |
| Previous contribution |  |  | 0.00<br>(0.13) | -0.05<br>(0.12) | -0.06<br>(0.12) |
| 2014 | 0.54*<br>(0.27) | 0.77*<br>(0.33) | 1.00**<br>(0.34) | 1.14**<br>(0.39) | 0.74*<br>(0.32) |
| 2016 | 0.71<br>(0.38) | 1.24***<br>(0.34) | 1.12<br>(0.62) | 1.77***<br>(0.37) | 1.25***<br>(0.33) |
| Male |  | 0.20<br>(0.33) |  | 0.31<br>(0.32) | 0.23<br>(0.33) |
| Age |  | 0.00<br>(0.02) |  | 0.01<br>(0.02) | 0.00<br>(0.02) |
| Married |  | 0.32<br>(0.53) |  | 0.55<br>(0.49) | 0.34<br>(0.54) |
| Number of living children |  | -0.04<br>(0.05) |  | -0.07<br>(0.05) | -0.04<br>(0.05) |
| Exposure to market |  | -0.07<br>(0.33) |  | -0.15<br>(0.42) | -0.08<br>(0.33) |
| Number of current coresidents |  | -0.05*<br>(0.02) |  | -0.08**<br>(0.02) | -0.05*<br>(0.02) |

*Note.* To establish robustness, we conducted the OLS regressions using an order logit regression. See STAR method for details. Values are unstandardized logit regression coefficients with standard errors in parentheses. All analyses are restricted to contributions in 2013, 2014, and 2016, and only to individuals with a previous contribution in the sample year prior.
\* $p < 0.05$ , \*\* $p < 0.01$ , \*\*\* $p < 0.001$

**Table S6.** OLS Regression of Current PG Contributions on Previous Play. Related to Campmates Influence Cooperative Behavior

|  | *b* | *SE* | *t* | *p* |
| --- | --- | --- | --- | --- |
| 2014 | 0.27 | 0.09 | 3.12 | 0.002 |
| 2016 | 0.18 | 0.11 | 1.60 | 0.111 |
| Cooperated<br>previously | 0.06 | 0.13 | 0.61 | 0.640 |
| Defected<br>previously | 0.09 | 0.15 | 0.61 | 0.540 |
| Camp mean | 0.72 | 0.06 | 12.00 | < 0.001 |

*Note.* Analysis regressed current PG contributions on participants' behavior in previous sample. Individuals who contributed as much or more than their previous campmates were coded as previous cooperators, whereas individuals who contributed less than their previous campmates were coded as previous defectors. Thus, the analysis compared previous behavior to individuals who did not participate in the previous sample. Robust standard errors were clustered on the individual and camp.

